# fMRI BOLD signals in the left angular gyrus and hippocampus are associated with memory precision

**DOI:** 10.1101/2025.06.10.658942

**Authors:** Mingzhu Hou, Paul F. Hill, Luke R. Pezanko, Ayse N. Z. Aktas, Arne D. Ekstrom, Michael D. Rugg

## Abstract

Prior functional magnetic resonance (fMRI) studies examining the neural correlates of retrieval success and precision have reported inconsistent results. Here, we examined the neural correlates of success and precision in a test of memory for spatial location. The present study extended prior findings by employing an experimental design that minimized temporal overlap between mnemonic and visuomotor processing. At study, participants viewed a series of object images, each placed at a randomly selected location on an imaginary circle. At test, studied images were intermixed with new images and presented to the participants. The requirement was to make a covert recognition memory judgment to each image and to attempt to recall its studied location, guessing if necessary. A cue signaling the requirement to make a location memory judgment was presented 4 seconds after image onset. Memory precision was quantified as the angular difference between the studied location and the location selected by the participant. In an analysis that combined the data from the present study and a closely similar prior study, we replicated prior reports that fMRI BOLD activity in the left angular gyrus (AG) and the hippocampus tracks memory precision on a trial-wise basis. Linear mixed effects modeling indicated that the activity in the two regions explained independent sources of variability in these judgments. In addition, multivoxel pattern similarity analysis revealed robust evidence for an item-level reinstatement effect (as indexed by encoding-retrieval overlap) in the left AG that was restricted to items associated with high precision judgments. These findings suggest that the hippocampus and the left AG play non-redundant roles in the retrieval and behavioral expression of high precision episodic memories.

## 1. Introduction

Episodic memory refers to consciously accessible memory for personally experienced, unique events (Tulving, 1983). Recent behavioral studies employing continuous memory metrics suggest that episodic memory retrieval can be dissociated into two components: retrieval success (the accessibility of the memory) and precision (the level of correspondence between the originally experienced event and the remembered event) (Gellerson et al., 2024; Harlow and Yonelinas, 2016; Korkki et al., 2020; Sutterer & Awh, 2016). As we discuss, below, whether retrieval success and precision engage common or dissociable neural processes is unclear.

Studies examining success and precision have typically employed ‘positional response accuracy’ tasks (Harlow and Donaldson, 2013). In these tasks, memory performance is assessed by a continuous metric that indexes the degree of correspondence between a feature of the study event and subsequent memory of that feature. For example, in our recent study (Hou et al., 2025) assessing memory precision for spatial location, participants studied a series of object images that were presented at different locations around a circle. In the subsequent test phase, both studied and new objects were presented at the center of the screen.

Participants were instructed to manually control a cursor and move it until it overlapped with the studied location of the object, guessing if necessary (see Figure 1: Exp1 test). They then signaled whether the object was studied or unstudied. Thus, as is typical of such tasks, participant responses provided a continuous metric of memory accuracy (precision) in the form of the angular difference between the judged and the studied location of correctly recognized objects. The distribution of these differences can be fit to a mixture model comprising a rectangular distribution that models the probability of random guess (i.e. unsuccessful retrieval), and a circular Gaussian distribution that models the probability of successful retrieval with varying precision.

Using functional magnetic resonance imaging (fMRI), four studies have examined the neural correlates of memory precision and success (Cooper et al., 2017; Hou et al., 2025; Korkki et al., 2023; Richter et al., 2016). Each study employed a positional response accuracy task and took a region of interest (ROI) approach to examine neural activity in the left angular gyrus (AG) and hippocampus. The results are mixed: Richter et al. (2016) and Cooper et al. (2017) reported that retrieval success was associated with enhancement of BOLD activity in the hippocampus, whereas memory precision was indexed by BOLD activity within the AG. By contrast, Korkki and colleagues (2023) reported that activity in both regions was sensitive to success and precision. In Hou et al. (2025), AG, but not hippocampal, activity scaled with precision, and we were unable to identify a neural correlate of success in either region.

Additionally, Hou et al. identified an item-related reinstatement effect in the AG that was specific for memories retrieved with high precision. That is, there was a reliable overlap in the patterns of activity elicited in this region between the study and the test trials that included these items.

In addition to the ROI analyses described above, Hou et al. (2025) also compared test trials associated with different classes of memory judgment [successful location retrieval, guesses (unsuccessful location retrieval) and correct rejections of unstudied objects] at the whole brain level. Robust judgment effects were identified in a variety of cortical and subcortical regions, including regions that have previously been implicated in visuomotor function, such as the dorsal parietal cortex (DPC), premotor cortex and the cerebellum. The effects primarily took the form of differences in the activity elicited by the unstudied objects relative to the other trial types. However, the task design employed by Hou et al. (2025; see Figure 1B) meant that location recall and the associated location judgment temporally overlapped. Thus, it was not possible to determine which, if any, of the identified effects were reflections of mnemonic processing as opposed to differences in the visuomotor demands associated with the different classes of judgment.

Here, we aimed to further investigate the neural correlates of retrieval success and precision. We employed an experimental design similar to that employed by Hou et al. (2025) but, crucially, we modified the test phase to incorporate a covert memory recall phase prior to the requirement to make the memory judgment, thereby temporally separating (and, we hoped, unconfounding) memory retrieval from the visuomotor demands of the associated judgment.

Thus, we were able to determine the extent to which the prior whole brain findings reported by Hou et al. (2025) and described above were a consequence of the demands of the continuous performance task rather than reflections of mnemonic processing. Additionally, and as described below, we were also able to combine the data from the present and prior experiments to examine the neural correlates of precision and success with greater statistical power than that afforded by the individual experiments. To differentiate the two experiments, the experiment described by Hou et al. (2025) is referred to below as experiment 1, and the current study is referred to as experiment 2.

## 2. Methods

The methods employed in experiment 1 have been described in detail previously (Hou et al., 2025). Here, we focus on experiment 2 and briefly describe the differences between the experiments. As detailed below, to maximize the comparability of the estimated neural signals, we used the same boxcar duration to model the neural activity in each experiment. Equivalent definitions of location hits, as well as those for ‘high’ and ‘low’ precision trials were employed across experiments for the same reason. Note that the results were essentially the same when experiment-specific boxcar durations and precision-based definitions were employed.

### 2.1 Participants

Participants in experiment 2 comprised 24 young adults (mean age = 23 yrs, age range = 18-33 yrs, 11 female). A non-overlapping sample of 23 young adults participated in experiment 1. All participants were cognitively healthy, right-handed, had normal or corrected-to-normal vision, no history of neurological or psychiatric illness, and were not taking prescription medications that affected the central nervous system.

Informed consent was obtained in accordance with the University of Texas at Dallas Institutional Review Board guidelines. Participants were compensated at the rate of $30 an hour. For experiment 2, data from two additional participants were excluded due to near-chance levels of recognition memory.

### 2.2 Experimental items

The same set of experimental items, which comprised 136 images of everyday objects (see Figure 1) was employed in both experiments. 102 of the images were employed as study items and an additional 34 images were employed as new (unstudied) test items.

### 2.3 Procedure

The study and test tasks employed in each experiment are schematized in Figure 1. As is evident from the figure, there was a single study block followed by two consecutive test blocks in experiment 1, and by four consecutive test blocks in experiment 2.

At study, participants viewed 102 object images, each presented for 6 s at a random location on a virtual circle (radius = 4.9 deg). A small green cursor was also presented on the circle at least 60 deg distant from the image. Participants were instructed to use either the left or right button on a button box to move the cursor around the circle until it occupied the center of the image (see Figure 1), and to remember both the image and its location.

At test, all studied images were intermixed with 34 unstudied images. The ordering of the test images was pseudo-randomized so that no more than 3 old or new images occurred in succession. In experiment 2, the test image was presented for 1 s, followed by a 3-s presentation of a white circle with the same radius as the virtual circle employed during the study phase. Participants were instructed to recall the studied location of the image. They were then to move a cursor that was randomly located on the circle to the recalled location, again by using a button box, and guessing if necessary. After moving the cursor, participants signaled whether the presented object had been studied or unstudied. As can be seen from Figure 1, the test procedure in experiment 1 differed, in that there was no delay between the presentation of the test image and the requirement to use the cursor to indicate its studied location.

The experimental display was viewed via a mirror mounted on the scanner head coil that reflected a back projected image (viewing distance 102 cm). Participants pressed the buttons under their right index and middle fingers to move the cursor along the circle in both the study and test phases, and, at test, to respond ‘old’ or ‘new’ to the object image. The mappings between the button presses and the direction of the cursor movement and, separately, the old/new judgments, were counterbalanced across participants in each experiment.

In both experiments a 30-s break occurred in the middle of each study and test block, when a ‘rest’ cue was presented at the center of the screen. Participants were instructed to relax during the breaks until the cue disappeared.

**Figure 1.**
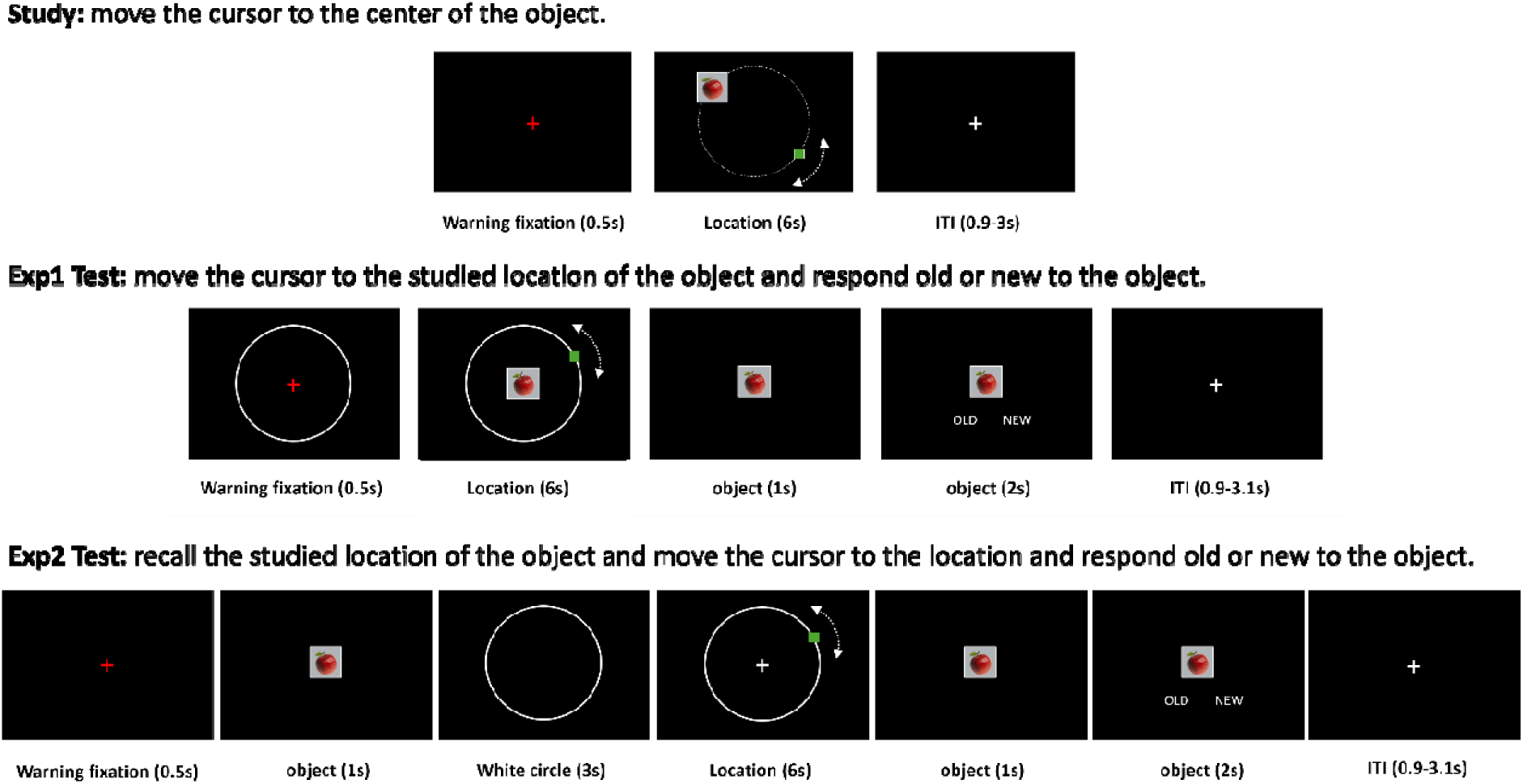
Schematic depiction of the study and test tasks for experiments 1 and 2. ITI: inter-trial interval.

### 2.4 MRI acquisition and preprocessing

MRI acquisition and preprocessing methods were identical across the two experiments. The data were acquired with a Siemens PRISMA 3T MR scanner equipped with a 32-channel head coil. Functional scans were acquired with a T2□-weighted echoplanar sequence [TR 1560 ms, TE 30 ms, flip angle 70°, field-of-view (FOV) 220 mm, multiband factor = 2, 48 slices, voxel size 2.5 x 2.5 x 2.5 mm, 0.5 mm inter-slice gap, anterior-to-posterior phase encoding direction]. A T1-weighted image was acquired with an MP-RAGE pulse sequence (TR = 2300 ms, TE = 2.41 ms, FOV 256 mm, voxel size = 1 x 1 x 1 mm, 160 slices, sagittal acquisition). A field map was acquired after the functional scans using a double-echo gradient echo sequence (TE 1/2 = 4.92□ms/7.38 ms, TR = 520□ms, flip angle = 60°, FOV 220□mm, 48 slices, 2.5mm slice thickness). Eye movement data were collected from both the study and retrieval tasks in experiment 2. They will be reported in a separate publication.

Data preprocessing was performed with the SPM12 software package (Wellcome Department of Imaging Neuroscience, London, UK: www.fil.ion.ucl.ac.uk/spm) implemented in MATLAB R2018b (experiment 1) and MATLAB 2023a (experiment 2). The functional volumes were field-map corrected, realigned, reorientated to the anterior commissure – posterior commissure line and spatially normalized to SPM’s MNI EPI template. The normalized images were resampled to 2.5 mm isotropic voxels and smoothed with a 6 mm Gaussian kernel. Before they were entered into the participant-wise GLMs the functional data from the different test blocks were concatenated using the spm_concatenate.m function. Anatomical images were normalized to SPM’s MNI T1 template.

### 2.5 Behavioral analyses

#### 2.5.1 Item recognition performance

Item memory (Pr) was quantified as the proportion of correctly recognized old items (item hits) minus the proportion of new items that were incorrectly endorsed as old (false alarms).

#### 2.5.2 Retrieval success and precision

Measures of retrieval success and precision were derived from the distributions of distance error (the difference between judged and actual study location in deg) of item hits. To enable comparability with prior findings, the distributions were fit to a two-component mixture model using the standard mixture modeling procedure implemented in the MemToolbox (Suchow et al., 2013). The model estimated a rectangular distribution reflecting the proportion of ‘guess’ responses (g), and a von Mises distribution that captured accurate location memory judgments of varying precision [characterized by the concentration parameter (Kappa) of the distribution]. Participant-wise g and Kappa metrics were estimated with the FitMultipleSubjects_MLE() function in MemToolbox. As in prior studies (e.g. Hou et al., 2025; Korkki et al., 2023; Richter et al., 2016), we estimated the retrieval success rate (pT) as 1 – g. In addition to Kappa, we also calculated a model-independent metric of memory precision, the mean distance error of item hit trials for each participant (Gellerson et al., 2024; Nilakantan et al., 2018). Unlike Kappa, this metric is meaningful at the single-trial level, and was employed in the trial-wise fMRI analyses reported below.

For each experiment, across-participant distance errors were fit to the mixture model using the MLE() function. Item hit trials were divided into successful (location hit) and guess trials depending on whether the associated distance errors had a < 5% chance of fitting the group-wise von Mises distribution as estimated with the “vonmisecdf” function from www.paulbays.com/toolbox/.

### 2.6 fMRI analyses

#### 2.6.1 ROI-based trial type analyses

Using a whole brain analysis based on an F contrast among location hits, guesses and correct rejections we identified robust effects of trial type in several cortical regions in our original report (Hou et al., 2025). As was mentioned in the introduction, the identified effects might reflect the varying visuomotor demands that were associated with the different trial types. In experiment 2, we sought to reduce these demands during memory retrieval by including a covert memory recall phase prior to the memory judgment phase (see Figure 1). To examine whether this procedural modification influenced the whole brain effects identified in Hou et al. (2025), we constructed participant-level GLMs analogous to those described in the original report. For the participant-wise GLMs, a 4 s boxcar function, which onset concurrently with the onset of the test item, was used to model neural activity synchronized to item onset in both experiments. Three event types of interest were included: location hits (item hit trials with absolute distance error < 47°, the model-based cutoff for the location hit based on data from experiment 1), guesses (item hit trials with absolute distance error > 46°), and correct rejections (CRs; correctly categorized unstudied items). For the sake of comparability with the whole brain analysis approach employed in Hou et al. (2025), a trial-specific measure of memory precision was included as a parametric modulator (although not reported here, as it is redundant with the linear mixed effects analyses reported below, the results derived from the parametric modulation model were highly similar to those reported previously). Other events, including false alarms, misses, and trials with absent or multiple old/new responses, were modeled as events of no interest. The GLMs also included as covariates six regressors modeling motion-related variance (three for rigid-body translation and three for rotation) and two (for experiment 1) or four (for experiment 2) constants for means across test blocks. Data from volumes with a transient displacement (relative to the prior volume) of >1 mm or >1° in any direction were modeled as covariates of no interest.

Parameter estimates were extracted from voxels within a 5 mm radius of the peak of each cluster identified in Hou et al. (2025) and subjected to a 3 (trial type) x 8 (region) x 2 (experiment) mixed effects ANOVA. The MNI coordinates of each peak were listed in Table 1.

**Table 1.** Regions demonstrating significant trial type effects in the whole brain analysis in Hou et al. (2025).

| Region | MNI |  |  |
| --- | --- | --- | --- |
|  | x | y | z |
| Left dorsal parietal cortex (DPC) | -26 | -62 | 52 |
| Right DPC | 14 | -72 | 52 |
| Left premotor cortex | -23 | -2 | 52 |
| Right premotor cortex | 30 | 3 | 55 |
| Left dorsal prefrontal cortex (DLPFC) | -56 | 26 | 32 |
| Left AG | -56 | -70 | 32 |
| Right AG | 62 | -52 | 38 |
| Left cerebellum | -20 | -84 | -38 |

#### 2.6.2 ROI-based categorical precision analyses

We conducted targeted univariate and multi-voxel pattern similarity analyses to examine the neural correlates of retrieval success and precision, focusing on fMRI signals in the left AG and the hippocampus. We employed a combined region of interest (ROI) in the left AG based on the peak precision effects reported by Korkki et al. (2023) and Richter et al. (2016) (MNI coordinates: -54, -66, 33 and -54, -54, 33); this is the same left AG ROI as that employed by Hou et al. (2025)]. The anatomically defined left and right hippocampi served as the hippocampal ROIs. These took the form of bilateral hippocampal masks that had been manually traced on the group mean T1 images averaged across the two experiments.

As in Hou et al. (2025), we contrasted the BOLD activity elicited by item hit trials according to level of memory precision. Three event types were modeled: high-precision hits (item hit trials with absolute distance error < 16 deg), low-precision hits (item hit trials with absolute distance error ranging from 25-40 deg) and guesses (item hit trials with absolute distance error > 59 deg). Mean trial numbers for high-precision hits, low-precision hits and guesses were 23 (range = 5-52), 11 (4-20) and 26 (5-43) for experiment 1, and 34 (13-61), 10 (2-17) and 20 (7-35) for experiment 2. Note that the results were unchanged after excluding data from participants with fewer than 5 low-precision hit trials. We employed an equivalent accuracy range in each experiment (15 deg) when defining the high and low-precision trials to reduce the potential influence of differential accuracy ranges on the neural estimates associated with the two trial types. The 10-deg boundary between the high vs low-precision trials was inserted to enhance possible categorical differences between trial types. To minimize the likelihood of including trials that more properly belonged to the location hit distribution, guess trials were defined as those where the distance error was substantially below the cut-off. All other events were modeled as events of no interest.

Parameter estimates associated with the different trial types were extracted from the predefined AG and hippocampal ROIs. They were subjected to separate repeated measures ANOVAs employing the factors of trial type (high precision, low precision, guess) and experiment. The factor of hemisphere was also included in the ANOVA of the activity elicited in the hippocampal ROIs. Precision effects were operationalized as greater activity for high-precision than low-precision trials, while retrieval success effects were defined as greater activity for low-precision trials than guesses.

#### 2.6.3 Additional categorical precision analyses in the hippocampus

We conducted additional exploratory analyses to further examine the role of the hippocampus in memory precision and success. Participant-wise parameter estimates derived from the first level GLMs described in the preceding section were subjected to a mixed effects ANOVA (factors of trial type and experiment) implemented in SPM12. To identify voxels where activity varied across the three trial types (high precision hits, low precision hits, guesses) in an unbiased manner, we employed an F contrast with thresholds of p < 0.05 and 20 contiguous voxels (see, Bakker et al., 2008 and Lacy et al., 2011 for examples of a similar approach). The contrast, which leaves the profile of response magnitudes across the trial types free to vary, was restricted to voxels that fell within the bilateral hippocampal mask. Participant-wise mean across-voxel parameter estimates were extracted from each cluster identified with the contrast and entered into a 3 (trial type) x 2 (experiment) mixed ANOVA.

#### 2.6.4 Linear mixed effects analyses

We constructed a set of linear mixed effects (LME) regression models to examine possible associations between BOLD activity in the left AG and hippocampus and the trial-wise absolute distance error of location hits. We also employed LMEs to ascertain whether any identified associations were independent of each other. We employed across-voxel mean parameter estimates derived from the AG ROI and the functionally defined hippocampal cluster (see preceding section) as predictors of absolute distance error. The models followed the general format:

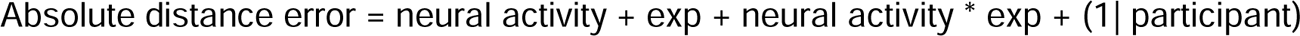

where absolute distance error refers to the absolute distance between the estimated and actual studied location in deg (0-46 deg), neural activity refers to retrieval-related BOLD signals in either left AG or the hippocampus, and Exp refers to experiment (experiment 1 coded as 0, experiment 2 coded as 1). Participants were modeled as a random effect with a variable intercept. Because a singular fit error indicative of model overfitting was evident in most of the initial models, random slopes for the variable of neural activity were not specified.

#### 2.6.5 Multi-voxel pattern similarity analyses

We conducted multi-voxel pattern similarity analyses (PSA) to examine whether memory precision was associated with the level of correspondence between item-specific patterns of neural activity elicited during encoding and retrieval. In each experiment, participant-wise single-trial beta estimates from the study and test phases were derived from first-level GLMs employing the least squares all (LSA) approach (Abdulrahman and Henson, 2016; Mumford et al., 2012). Trial wise neural activity elicited by the study items was modeled with a 6 s duration boxcar. Each study event was modeled as a separate event of interest, and the 30 s rest period that occurred midway through the study block, the six motion regressors, and a constant modeling the mean BOLD signal in the block were included as covariates of no interest. An analogous single-trial GLM was employed for the test blocks, with event-related neural activity modelled as a 4 s boxcar.

We examined retrieval-related reinstatement effects associated with high-precision location hits, low-precision location hits and guess trials. For each trial belonging to each class of memory judgment, encoding- and retrieval-related trial-wise parameter estimates were extracted from all voxels falling within a given ROI. As in Hou et al. (2025) we employed three ROIs, namely, the left and right anatomically defined hippocampus and left angular gyrus [the ‘PGap’ parcel specified in the Anatomy Toolbox v3.0 (Eickhoff et al., 2005, 2006; 2007)]. Reinstatement for any given item was operationalized as the difference between within-item and across-item study-test similarity. Within-item similarity was calculated as the across-voxel Fisher-z transformed correlation between a given study trial and its corresponding test trial.

Across-item similarity was computed as the mean of the Fisher-z transformed correlations between the same study trial and all other test trials belonging to the same class of memory judgment. For each participant, the within minus between similarity estimates were averaged across all trials belonging to each class of judgment. One sample t-tests were employed to assess whether, across participants, the resulting similarity measures differed reliably from zero. Trial type (high-precision, low-precision, guess) x experiment mixed effects ANOVAs was conducted to examine associations between reinstatement strength and memory precision in each ROI.

## 3. Results

### 3.1 Behavioral results

The distributions of distance errors of all item hits pooled across participants from the two experiments are illustrated in Figure 2. The model-derived estimate of precision (Kappa) was 6.43 for experiment 1 and 11.04 for experiment 2. The retrieval success rate (pT) was 0.51 and 0.62 for experiments 1 and 2, respectively. The model-based cutoff for the location hit vs guess trials was estimated as +/-47 deg for experiment 1 and +/-35 deg for experiment 2.

Table 2 shows participant-wise memory performance in each experiment. Both the mean Kappa and the absolute distance error for item hits differed significantly between experiment 1 and 2, indicative of higher precision in experiment 2. Item recognition performance was comparable between the two experiments.

**Figure 2.**
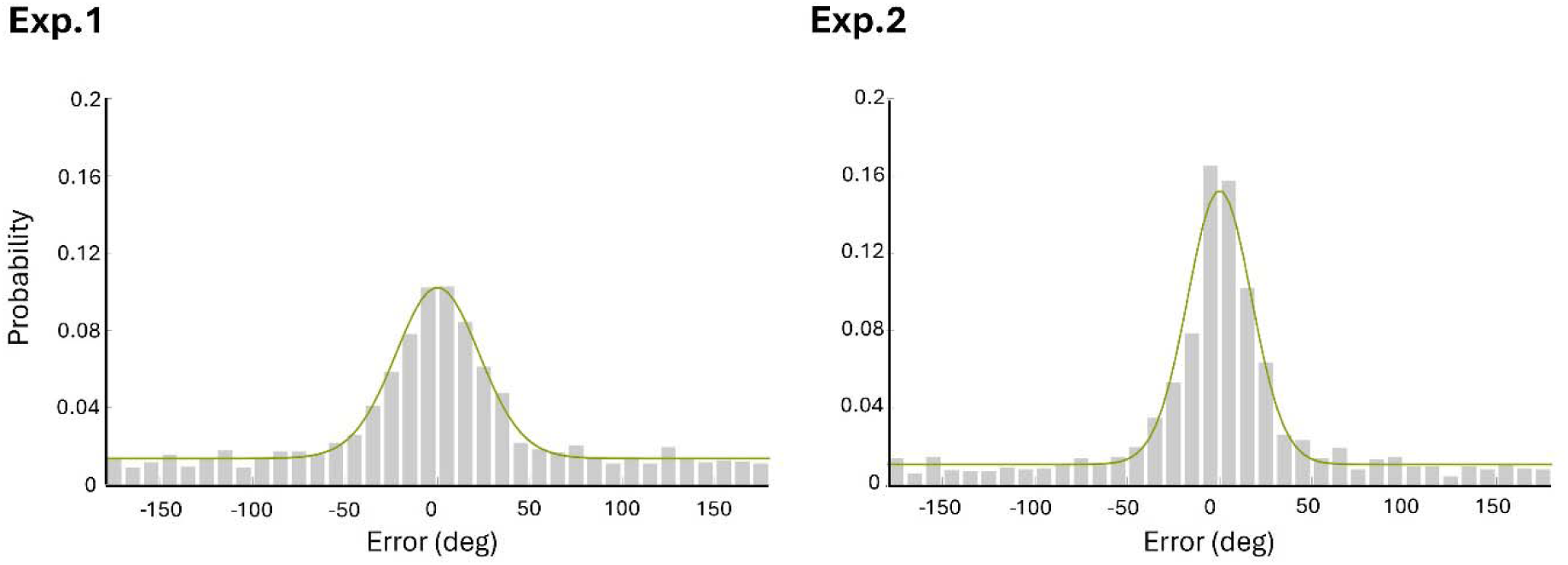
Distribution of distance errors for item hit trials across participants in experiment 1 and 2 (trial number = 1777 and 1917 for experiment 1 and 2, respectively). Each curve indicates the fit of the von Mises + uniform distributions mixture model.

**Table 2.** Mean participant-wise performance for the spatial precision task and item recognition.

|  | Exp 1 | Exp 2 | t | p |
| --- | --- | --- | --- | --- |
| Kappa | 5.38 (2.59) | 12.43 (6.37) | 5.01 | **< 0.001** |
| Guess rate | 0.43 (0.20) | 0.39 (0.17) | 0.82 | 0.415 |
| Absolute distance error | 53.99 (15.39) | 42.16 (12.72) | 2.88 | **0.006** |
| Item hit rate | 0.76 (0.11) | 0.78 (0.13) | 0.74 | 0.465 |
| Correct rejection rate | 0.81 (0.13) | 0.86 (0.12) | 1.23 | 0.224 |
| Pr | 0.60 (0.19) | 0.65 (0.17) | 1.14 | 0.259 |

### 3.2 fMRI results

#### 3.2.1 ROI-based trial type analyses

We also examined the effects of trial type in the regions originally reported by Hou et al. (2025). Results of the ANOVA contrasting the parameter estimates for location hits, guesses and CRs are shown in Table 3. As is evident from the table, there was a significant trial type x region x experiment interaction, indicating that the effect of trial type varied with both region and experiment. To unpack the interaction, we conducted pairwise t-tests among trial types in each region and experiment. As is illustrated in Figure 3, unlike in experiment 1, in experiment 2 none of the contrasts between trial types were significant in the left AG, cerebellum, DLPFC and right AG. However, effects in bilateral premotor cortex and DPC were significant in both experiments, taking the form of higher parameter estimates for location hits and guesses than for CRs.

**Table 3.** Results of ANOVAs comparing location hits, guesses and CR trials in the regions identified by Hou et al. (2025).

|  | Results |
| --- | --- |
| Trial type | **F(1.90, 85.66) = 7.75, p < 0.001, partial $\eta^2$ = 0.147** |
| Region | **F(3.32, 149.33) = 92.99, p < 0.001, partial $\eta^2$ = 0.674** |
| Experiment | F(1, 45) = 1.72, p = 0.196, partial $\eta^2$ = 0.037 |
| Trial type x Region | **F(5.29, 237.97) = 27.26, p < 0.001, partial $\eta^2$ = 0.377** |
| Trial type x Experiment | F(1.90, 85.66) = 0.46, p = 0.624, partial $\eta^2$ = 0.010 |
| Region x Experiment | **F(3.32, 149.33) = 7.33, p < 0.001, partial $\eta^2$ = 0.140** |
| Trial type x Region x Experiment | **F(5.29, 237.97) = 6.10, p < 0.001, partial $\eta^2$ = 0.119** |

**Figure 3.**
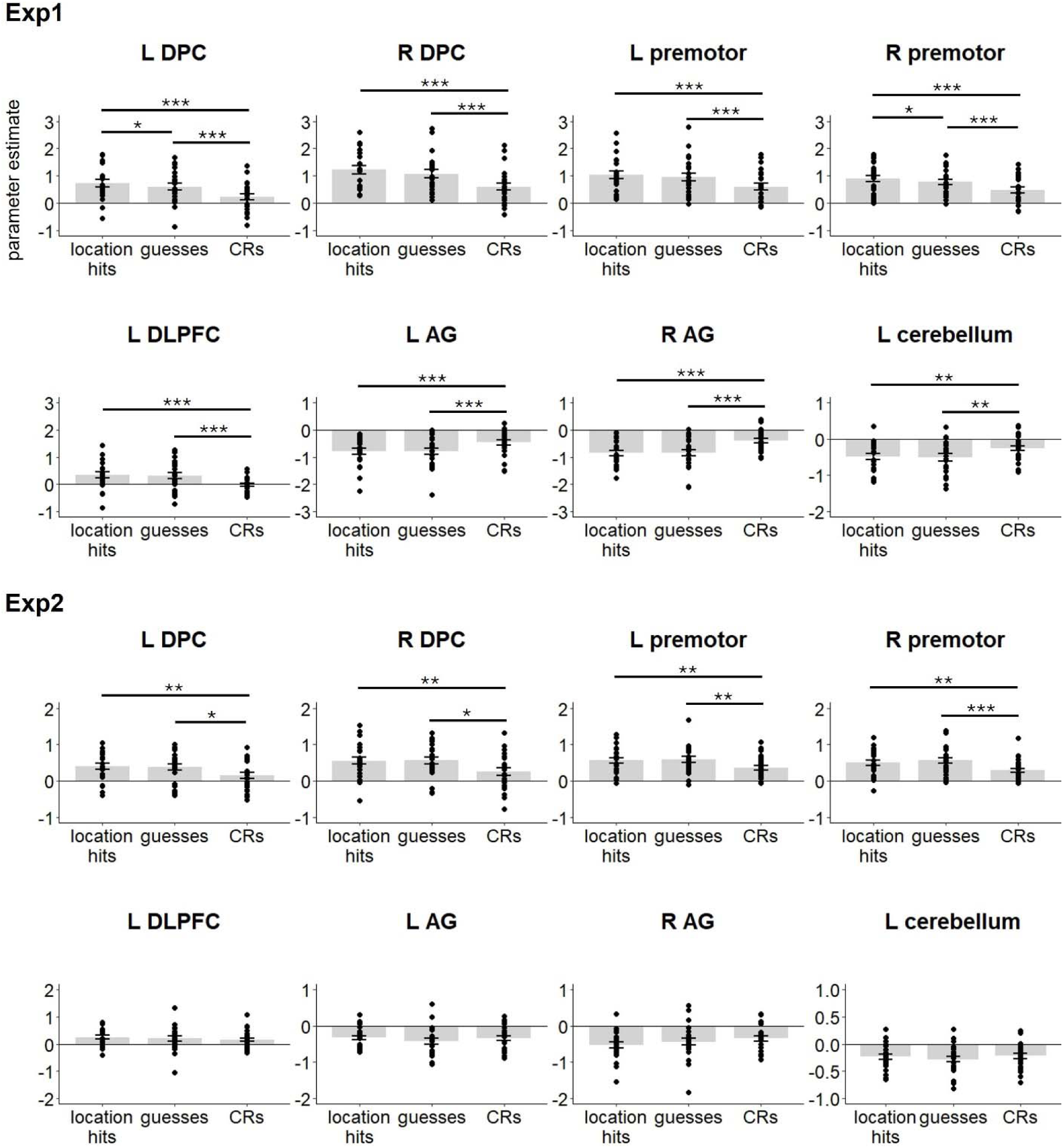
Parameter estimates extracted from the cortical regions identified by the whole brain analyses in Hou et al. (2025). Error bars indicate standard errors of the mean. * p < 0.05, ** p < 0.01, *** p < 0.001.

#### 3.2.2 Categorical precision analyses

The results of the ANOVAs contrasting the parameter estimates associated with high precision, low precision and guess trials are listed in Table 4 separately for the AG and hippocampal ROIs. In the case of the AG, the main effect of trial type was significant. Follow-up analyses revealed that activity associated with high-precision hits was greater than that for low precision hits (t46 = 3.01, p =0.004, Cohen’s d = 0.44, see Figure 4). There were no other significant differences (ps > 0.077). As is evident from Table 4, the ANOVA of the data from the AG also revealed a significant main effect of experiment, indicative of greater overall BOLD activity in experiment 2 (M = -0.42) than experiment 1 (M = -0.81). Turning to the hippocampus, there was no effect of trial type, while the main effect of experiment was again reliable (experiment 1: M = -0.17; experiment 2: M = -0.07). In summary, we were able to identify memory precision effects in the AG but not in the hippocampus. No effects of retrieval success, defined as greater activity for low-precision hits than guesses, were evident in either ROI.

**Table 4.** Results of ANOVAs comparing high-precision, low-precision and guess trials in the a priori defined AG and the hippocampal ROIs.

| Region | Results |
| --- | --- |
| *Combined AG* |  |
| **Trial type** | **$F(1.99, 89.33) = 4.82$, $p = 0.010$, partial $\eta^2 = 0.097$** |
| **Experiment** | **$F(1, 45) = 10.78$, $p = 0.002$, partial $\eta^2 = 0.193$** |
| Trial type x Experiment | $F(1.99, 89.33) = 0.17$, $p = 0.840$, partial $\eta^2 = 0.004$ |
| *Hippocampus* |  |
| Trial type | $F(1.79, 80.56) = 1.33$, $p = 0.269$, partial $\eta^2 = 0.029$ |
| Hemisphere | $F(1, 45) = 2.65$, $p = 0.110$, partial $\eta^2 = 0.056$ |
| **Experiment** | **$F(1, 45) = 4.64$, $p = 0.037$, partial $\eta^2 = 0.093$** |
| Trial type x Hemisphere | $F(1.73, 78.03) = 1.61$, $p = 0.209$, partial $\eta^2 = 0.035$ |
| Trial type x Experiment | $F(1.79, 80.56) = 0.08$, $p = 0.908$, partial $\eta^2 = 0.002$ |
| Hemisphere x Experiment | $F(1, 45) = 0.03$, $p = 0.860$, partial $\eta^2 = 0.001$ |
| Trial type x Hemisphere x Experiment | $F(1.73, 78.03) = 0.11$, $p = 0.866$, partial $\eta^2 = 0.003$ |

**Figure 4.**
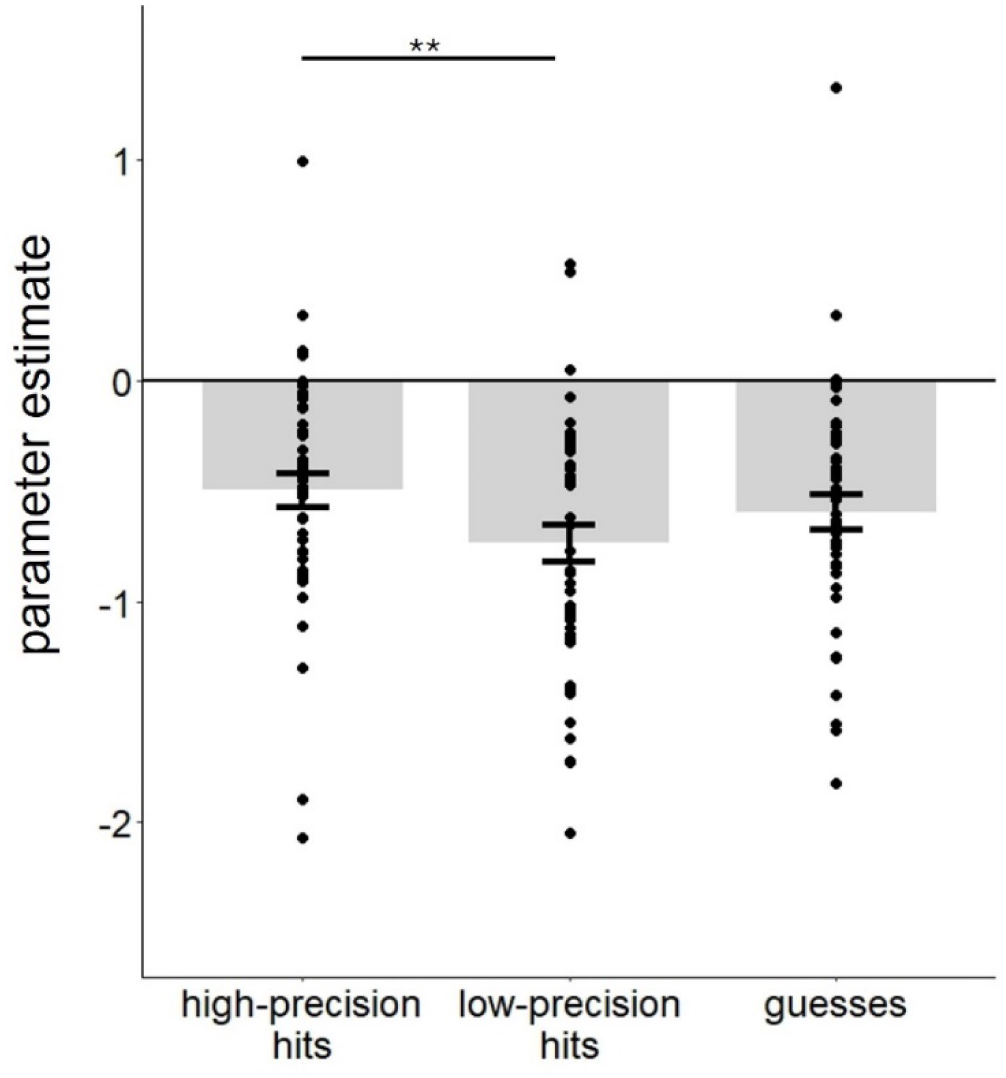
Parameter estimates of high-precision, low-precision and guess trials from the combined AG ROI. Error bars indicate standard errors of the mean. ** p < 0.01.

#### 3.2.3 Additional categorical precision analyses in the hippocampus

Employing the approach described in section 2.6.3, we identified a single cluster in the right hippocampus (peak 35, -20, -18, k = 41) that demonstrated a significant main effect of trial type. We conducted a 3 (trial type) x 2 (experiment) ANOVA on the parameter estimates extracted from the cluster, the results of which are shown in Table 5. Unsurprisingly, given how the cluster was identified, there was a main effect of trial type. Crucially, and undetermined by the main effect, follow-up pairwise t-tests indicated that the parameter estimates associated with high-precision trials were greater than those for either low-precision or guess trials (respectively, t46 = 3.33, p = 0.002, Cohen’s d = 0.49; t46 = 3.11, p = 0.003, Cohen’s d = 0.45). The latter two trial types did not significantly differ (p = 0.293, see Figure 5). Therefore, this analysis identified a significant precision effect in the right hippocampus. The main effect of experiment was also significant, reflecting greater activity in experiment 2 (M = - 0.01) than experiment 1 (M = -0.12).

**Table 5.**
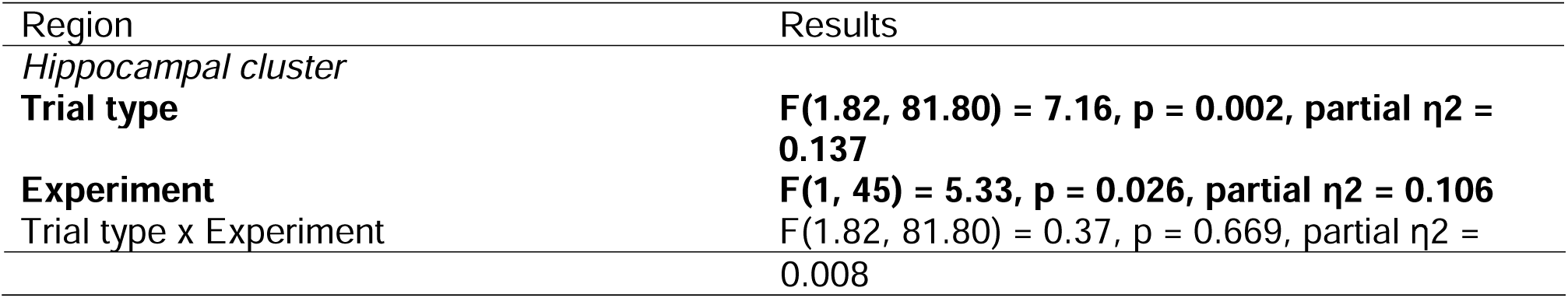
Results of ANOVAs comparing item hit trials associated with high, low precision and guess in the functionally defined hippocampal cluster.

**Figure 5.**
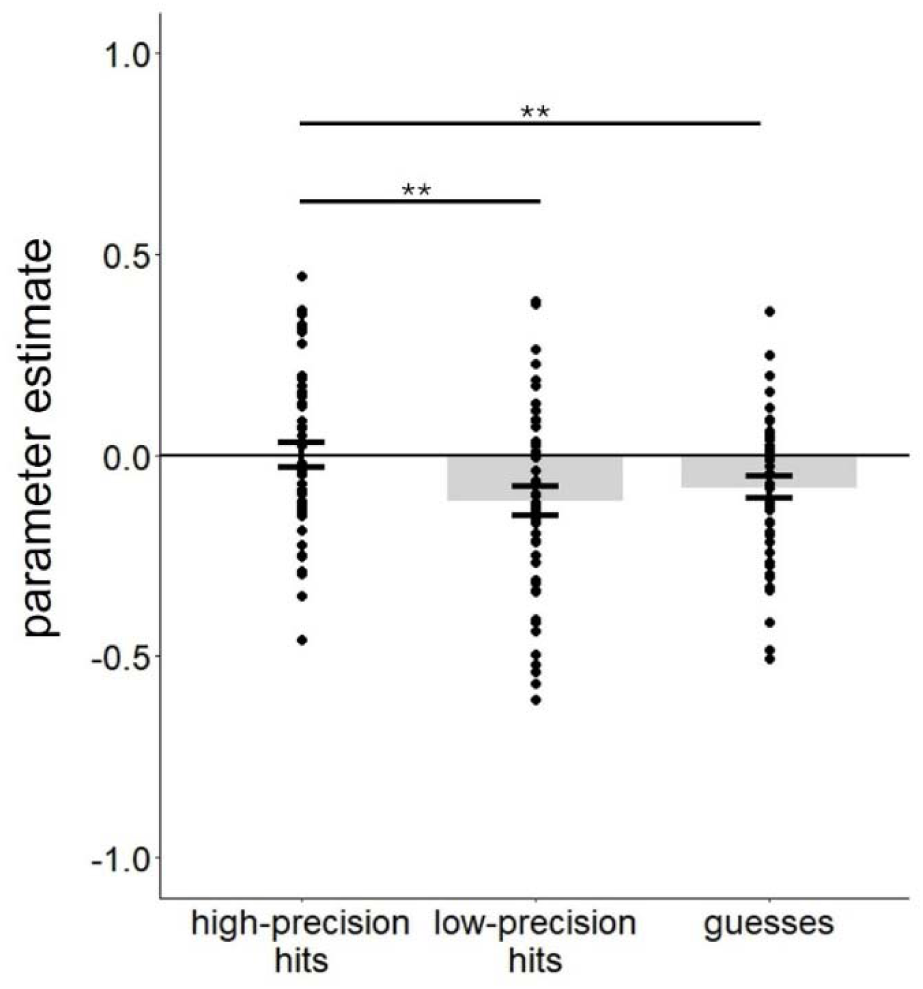
Parameter estimates associated with high-precision, low-precision and guess trials from the cluster identified by the ANOVA targeted on the hippocampus. Error bars indicate standard errors of the mean. ** p < 0.01.

#### 3.2.4 Linear mixed effects analyses

As described in the methods, we employed LME models to examine the relationships between trial-wise neural activity in the AG and hippocampus and memory precision. In these models, we employed parameter estimates extracted from the AG ROI and the functionally defined hippocampal cluster (see preceding section) as predictors of the absolute distance error of location hit trials. The results are summarized in Table 6. As is evident from the table, trial-wise BOLD activity in both the AG and hippocampus was predictive of absolute distance error, such that greater activity was associated with lower error (and thus higher precision, see Figure 6). Of importance, when they were included in the same regression model, hippocampal and AG parameter estimates independently predicted memory precision.

**Table 6.** Results from linear mixed models examining the relationships between memory precision and trial-wise BOLD activity extracted from the combined AG and the hippocampal cluster.

| Parameter | B(SE) | df | t | p |
| --- | --- | --- | --- | --- |
| *AG* |  |  |  |  |
| **AG** | **-0.51 (0.18)** | **2382.58** | **2.85** | **0.004** |
| **Exp** | **-3.15 (0.74)** | **47.43** | **4.27** | **< 0.001** |
| *Hippocampus* |  |  |  |  |
| **Hippocampus** | **-0.95 (0.33)** | **2439.80** | **2.89** | **0.004** |
| **Exp** | **-3.24 (0.74)** | **46.96** | **4.38** | **< 0.001** |
| *AG +*<br>*Hippocampus* |  |  |  |  |
| **Hippocampus** | **-0.74 (0.34)** | **2442.83** | **2.19** | **0.028** |
| **AG** | **-0.40 (0.19)** | **2401.01** | **2.13** | **0.033** |
| **Exp** | **-3.09 (0.74)** | **47.49** | **4.19** | **0.001** |

Note. In an initial iteration of these model, the neural activity x experiment term was not significant in the models involving either AG or hippocampal activity (ps > 0.524).

**Figure 6.**
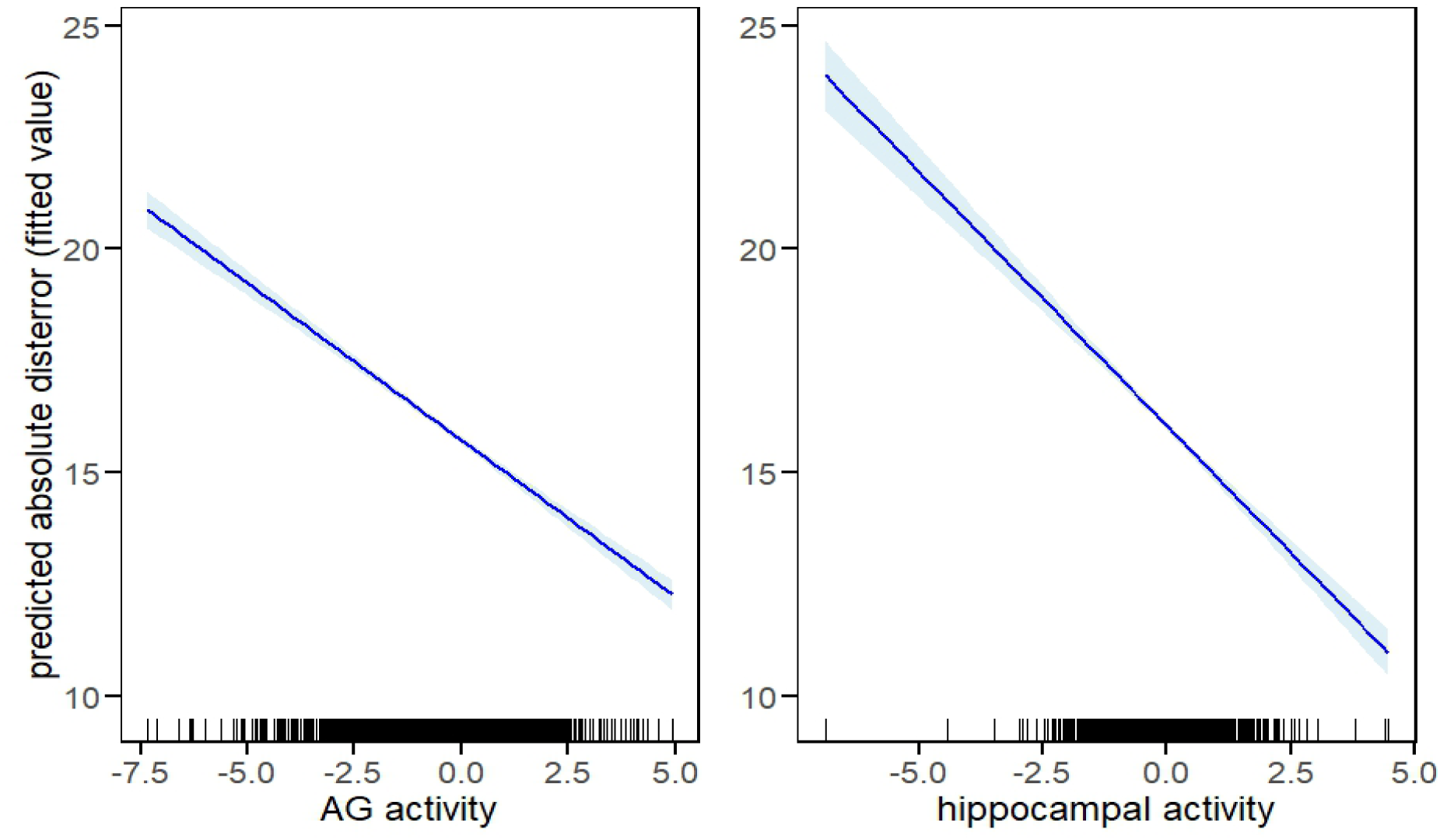
Plots depicting the relationships between trial-wise neural activity in the AG and the hippocampus and the model fitted values of memory precision for location hits, as derived from the LME model in which the two regions served as joint predictors of precision. The shaded areas reflect the 95% confidence intervals. The results were unaltered by the exclusion of trials on which fMRI BOLD parameter estimates were greater than 3 SDs from the mean for either the left AG or the hippocampus.

#### 3.2.5 Item-level reinstatement

Using PSA, we examined whether memory precision was associated with item-level reinstatement, namely the correspondence between item-specific patterns of neural activity elicited during encoding and retrieval. A reliable item-level reinstatement effect was evident in the anatomically defined AG (see Methods) in association with high-precision trials (t46 = 2.56, p = 0.014, Cohen’s d = 0.37, see Figure 7). By contrast, neither low precision trials nor guesses were associated with above-chance reinstatement (ps > 0.550). The 3 (trial type) x 2 (experiment) mixed ANOVA for the AG ROI revealed a significant main effect of experiment, indicative of greater mean pattern similarity in experiment 1 (M = 0.02) than experiment 2 (M = - 0.01). Although the main effect of trial type and the trial type x experiment interaction were both non-significant (ps > 0.067), planned pairwise t-tests revealed a greater reinstatement effect for high than for low precision trials (t46 = 2.42, p = 0.020, Cohen’s d = 0.35; for the comparison between high precision and guess trials, p = 0.289, see Figure 7).

We repeated the ANOVA after excluding the data from the participant with the outlying data point (> 3 SDs from the group mean) indicated in Figure 7. The ANOVA revealed significant main effects of trial type (F1.85, 81.52 = 5.01, p = 0.010, partial η2 = 0.102) and experiment (F1, 44 = 7.29, p = 0.010, partial η2 = 0.142), while the trial type x experiment interaction was not significant (p = 0.553). Pairwise contrasts indicated that the reinstatement effect associated with high precision hits was now greater than either low precision or guess trials (respectively: t45 = 2.82, p = 0.007, Cohen’s d = 0.42; t45 = 2.54, p = 0.015, Cohen’s d = 0.37) while the latter two trial types still did not differ significantly (p = 0.369). No reinstatement effects were identified for any trial type in the hippocampal ROIs (ps > 0.513).

**Figure 7.**
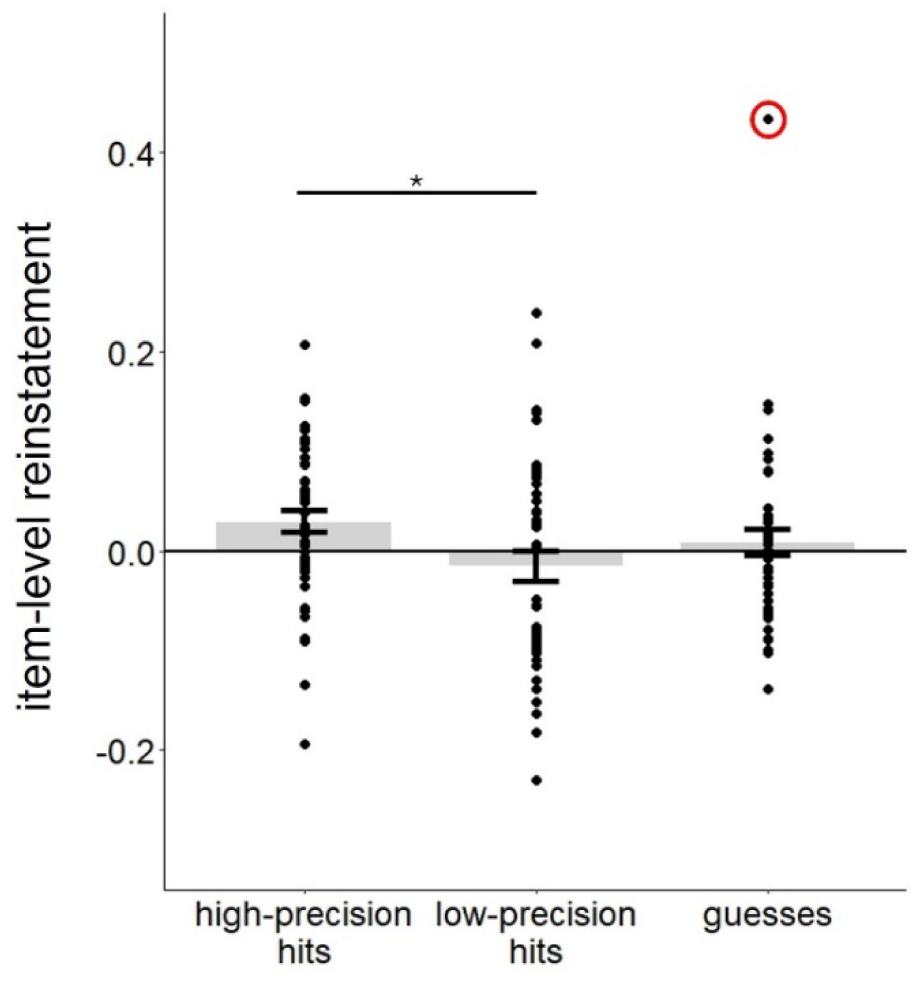
Estimates of item-level reinstatement in the anatomically defined AG. Excluding data from the participant with the outlying data point (circled), the estimate of high-precision hits was greater than both low-precision hits and guesses (ps < 0.016, see main text). * p < 0.05.

## 4. Discussion

The present study examined the neural correlates of retrieval success and memory precision for spatial location, building on the findings reported by Hou et al. (2025) by employing an experimental design that reduced the temporal overlap that previously existed between visuomotor and mnemonic processing. With the modified design, several of the cortical effects differing according to the type of memory judgment were no longer detectable. In analyses that extended across the prior and present experiment, we identified significant precision effects in the left AG and the hippocampus. Additionally, linear mixed effect analyses revealed that neural activity in these two regions independently predicted memory precision on a trial-wise basis.

Moreover, in the left AG, across-voxel patterns of neural activity elicited in the study and test phases demonstrated a reliable overlap for objects whose locations were remembered with high precision. These findings are discussed in more detail below.

An important goal of the current study was to elucidate the functional significance of the whole brain trial type effects reported by Hou et al. (2025). As was noted in the Introduction, Hou et al. identified significant differences in the neural activity associated with location hits, guesses, and correct rejections across a variety of cortical and subcortical regions. However, because the test design incorporated a temporal overlap between location recall and the corresponding location judgment, the functional significance of these differences was ambiguous. Here, we modified the design employed by Hou et al. (2025) by incorporating a covert memory recall phase prior to the memory judgment, hence removing the aforementioned overlap. With this modification, whole brain trial type effects were no longer detectable in the left DLPFC or the cerebellum. Thus, consistent with prior evidence implicating these regions in visuomotor function (for reviews, see Schall, 2015; Tzvi et al., 2022), these findings suggest that the judgment effects in these regions evident in experiment 1 were associated with the varying visuomotor demands associated with the different classes of trial [trial type effects were also absent in experiment 2 in the bilateral AG. We have no ready explanation why these effects, which took the form of enhanced activity for correct rejections (‘novelty effects’) should have been impacted by the change in experimental design]. By contrast, trial type effects remained significant in experiment 2 in dorsal parietal and premotor cortex, taking the form, as in experiment 1, of greater activity for location hits and guesses relative to correct rejections. We conjecture that these across-experiment effects reflect differences in either oculomotor activity or motor planning operations associated with the different classes of trial.

Given that we did not identify any reliable between-experiment differences in respect to the neural correlates of retrieval precision, we henceforth focus on the analyses of the combined dataset. Consistent with prior findings (Cooper et al., 2017; Korkki et al., 2023; Richter et al., 2016), we identified a memory precision effect in the left AG that took the form of greater BOLD activity for locations remembered with high relative to low precision. However, as was also reported for the analysis of experiment 1 alone (Hou et al., 2025; see also Cooper et al., 2017; Richter et al., 2016), we were unable to identify evidence of a retrieval success (low precision hits > guesses) effect in the left AG. On their face, these null findings appear to be inconsistent with the proposal that this region is sensitive to memory precision: precision obviously is greater for low precision judgments than for those made in the absence of any diagnostic information at all (guesses). As discussed previously (Hou et al., 2025), these null results might however reflect the fact that, while bereft of information diagnostic of location information, trials associated with ‘guesses’ were associated with recollection of non-diagnostic information. That is, while location hit trials were a consequence of the successful encoding of an item–location association, guess trials arose when participants devoted attention to a feature of the study event other than object location. Since memory content was not assessed for any feature apart from location in the present experiments, the validity of this proposal cannot be assessed based on current data. Future studies incorporating additional memory metrics into the positional response accuracy paradigm would help to test this possibility.

In addition to the findings for the left AG, we also identified a hippocampal cluster that exhibited a significant memory precision effect (we caution, however, that these findings were derived from an unplanned exploratory analysis, and should be treated as provisional). As was reviewed in the Introduction, out of the four published fMRI studies that examined the neural correlates of precision, only Korkki et al (2023) reported a significant precision effect in the hippocampus. This mixed evidence stands in contrast to lesion evidence indicating that patients with medial temporal lobe lesions that extend into the hippocampus demonstrate lower memory precision than patients with exclusively extrahippocampal lesions (see Introduction). The present findings are consistent with these prior results (and those of Korkki et al. 2023) in suggesting that the hippocampus plays a specific role in high fidelity recollection (see also Ekstrom & Yonelinas, 2020).

Of importance, our findings go beyond prior reports in demonstrating that fMRI BOLD activity in the left AG and hippocampus explain unique components of across-trial variance in memory precision. These regions have long been held to play different roles in memory retrieval. The hippocampus is considered crucial for the encoding of item-context associations and for supporting the retrieval-related reinstatement of neocortical representations of these associations (e.g. Rugg et al., 2015). By contrast, among other possible functions, the left AG has been proposed to support a multi-modal ‘episodic buffer’ that makes episodic content available to executive control processes (King et al., 2015, Rugg & King, 2018; Vilberg and Rugg, 2009, see also Humphreys et al., 2021). The present results suggest that, in cognitively healthy young adults, the contribution of the left AG to memory precision adds to that of the hippocampus. Of course, this does not mean that the AG plays an independent role in precision-based memory judgments. Indeed, based on the available evidence, a likely scenario is that the mnemonic information on which the AG operates depends upon hippocampally-mediated retrieval processing Rugg et al., 2015; Staresina & Wimber, 2019; Vilberg & Rugg, 2012).

In addition to the univariate precision effects discussed above, the left AG also demonstrated an item-related reinstatement effect that were uniquely associated with high precision judgments. Consistent with our results, prior studies employing multivoxel analysis approaches have also reported AG reinstatement effects for study events that were later retrieved with high-fidelity (Kuhl & Chun, 2014; Lee et al., 2019). These findings raise the possibility that the AG plays a role in the reactivation or reinstatement of the fine-grained representations of prior events. It remains to be determined whether these effects reflect the fidelity with which the events are represented at the time of encoding, the fidelity of the retrieved episodic representations, or some combination of the two (see Hill et al., 2021, for evidence favoring the first of these possibilities). In contrast to these findings for the left AG, we were unable to identify an analogous reinstatement effect in the hippocampus. This null finding should be treated with caution, however, given the relatively coarse spatial resolution of the fMRI methods employed in the Hou et al. (2025) and the current study (2.5 mm isotropic voxels).

## 5. Limitations

There are several limitations to the current study. First, we only assessed location memory, leaving it unclear whether the findings, particularly those related to item reinstatement, generalize to memory for other features. Second, we were unable to assess the potential influence of memory for nondiagnostic information on the BOLD responses elicited by studied test items. And third, as already alluded to, the spatial resolution of the fMRI data in the present study was too low to identify memory effects at the level of hippocampal subfields. These limitations are all addressable in future research.

## 6. Conclusion

Contrasts of the whole brain results from our prior (Hou et al., 2025) and the present study revealed that several, but not all of those results were likely a consequence of employing an experimental design in which mnemonic and visuomotor processing demands overlapped temporally. The functional significance of the results common to the two studies remains to be established. Additionally, analysis of the combined datasets revealed that, as indexed by fMRI BOLD signals, neural responses in the left angular gyrus and the hippocampus explain independent components of trial-wise variance in memory precision. These findings suggest that these regions play non-redundant roles in supporting the retrieval and behavioral expression of high-fidelity episodic memories.

## Acknowledgments

This work was supported by the National Institute of Neurological Disorders and Stroke (grant number R01NS114913). We thank our experimental participants for volunteering their time.

